# RNA polymerases from Low G+C Gram Positive Bacteria

**DOI:** 10.1101/2021.06.06.447298

**Authors:** Michael Miller, Aaron J. Oakley, Peter J. Lewis

## Abstract

The low G+C Gram positive bacteria represent some of the most medically and industrially important microorganisms. They are relied on for the production of food and dietary supplements, enzymes and antibiotics, as well as being responsible for the majority of nosocomial infections and serving as a reservoir for antibiotic resistance. Control of gene expression in this group is more highly studied than in any bacteria other than the Gram negative model *Escherichia coli*, yet until recently no structural information on RNA polymerase (RNAP) from this group was available. This review will summarise recent reports on the high resolution structure of RNAP from the model low G+C representative *Bacillus subtilis*, including the role of auxiliary subunits δ and ε, and outline approaches for the development of antimicrobials to target RNAP from this group.

## Introduction

RNAP structures from multiple bacterial species have been determined. Within molecular microbiology, two organisms have dominated research into transcription and its regulation. *Escherichia coli*, a Gram-negative bacterium, has been extensively studied to understand fundamental mechanistic aspects of transcription and its regulation *e.g*. (1). *Bacillus subtilis*, a Gram-positive organism, has been extensively studied for the regulatory processes associated with the initiation of differential gene expression followed by compartment-specific transcription activation and gene expression during the developmental process of sporulation (2).

At a structural level, RNAP from *E. coli* (RNAP_EC_) has been studied for many years and with the advent of modern cryo electron microscopy and single particle analysis techniques (cryoEM)(3), was one of the first pseudo-atomic resolution multi-subunit RNAP structures solved (4). Prior to the advent of these current resources, high resolution structural data from X-ray crystallography was largely obtained from thermophiles for which little molecular biology data on transcription and its regulation had been performed. Knowing the structure of RNAP_EC_ allowed the reconciliation of structural and functional data in one system which has enabled profound new insights into the mechanisms of transcription and its regulation *e.g*. (5,6)(7,8).

Such structural data on the Gram-positive *B. subtilis* system has been lagging, but high resolution structures of several important complexes of *B. subtilis* RNAP (RNAP_BS_) have been recently published that enable a similar reconciliation of structural and molecular data (9–11). Despite the considerable similarity between all multi-subunit RNAPs from bacteria, there are important mechanistic differences that can now be examined. For example, initiation complexes tend to undergo multiple rounds of abortive initiation prior to leaving the promoter region and entering the elongation phase in *E. coli*, but similar effects are not observed in *B. subtilis* (12,13). The concentration and identity of the initiating NTP ([iNTP]) also has a major effect on transcription efficiency in *B. subtilis* (14).

Increasing our understanding of transcription regulation through structure-function studies is particularly important as *B. subtilis* is an industrially significant organism used in the production of enzymes (proteases, lipases, amylases), surfactants, and antibiotics (bacitracin) which have highly complex regulatory circuits controlling the expression of their genes. As a member of the *Firmicutes*, it is closely related to many of the most important clinical pathogens such as *Staphylococcus, Streptococcus, Enterococcus*, and *Clostridia*. In this review we will examine the structure of RNAP_BS_ and compare it with that of other bacteria, focussing on its unique features and auxiliary subunits and include reference to homologous RNAPs from closely related *Firmicutes* pathogens and how this information could be exploited in structure-based drug design.

## Overall structure: RNA polymerases from the low G+C Firmicutes

All bacterial RNAPs have a similar overall subunit composition comprising two α subunits that form an asymmetric dimer scaffold upon which the catalytic β and β′ subunits assemble (Fig. 1). Due to the lack of lineage-specific inserts (see below) RNAP_BS_ is more compact (shorter and narrower) than other bacterial RNAPs: 150 Å × 112 Å × 123 Å (L × W × H), *vs* 157 × 153 × 136 Å; *E. coli*, 183 × 107 × 115 Å; *Mycobacterium smegmatis*, and 170.1 × 110.1 × 127.8 Å; *Thermus thermophilus* (10).

**Figure 1.**
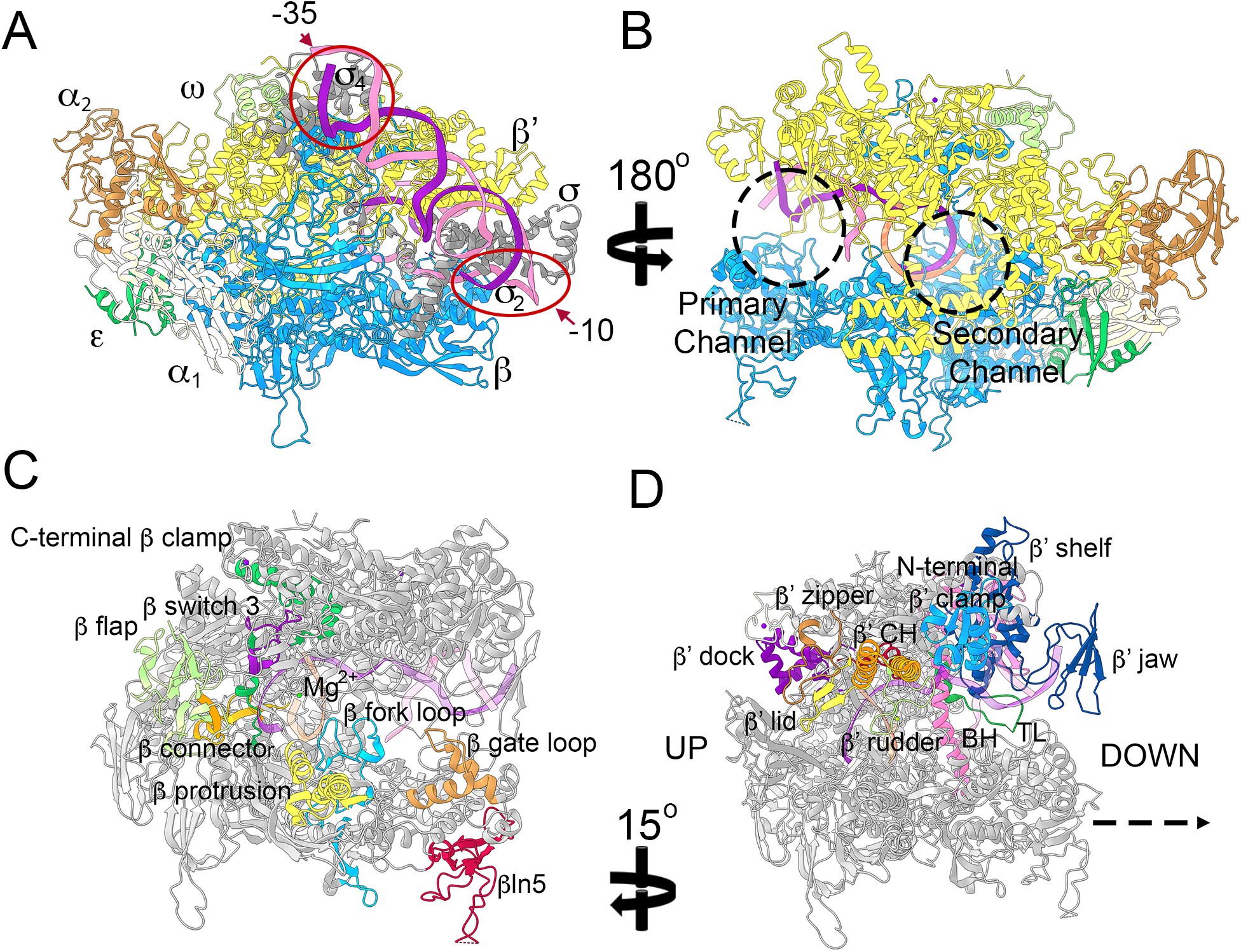
Structure of *B. sutbtilis* RNAP. Panel A shows the structure of RNAP holoenzyme, and Panel B an elongation complex. Subunit colouring; α_1_ cream, α_2_ brown, β blue, β′ yellow, ε dark green, ω pale green, σ grey. Template strand DNA is shown in dark purple, non-template strand DNA in pink, and RNA in orange. The −10 and −35 promoter elements are ringed in Panel A, and the primary and secondary channels circled in Panel B. Panels C and D show key functional elements of the β and β′ subunits, respectively, as defined by (11). β subunit elements in Panel B; C-terminal β clamp dark green, β switch 3 purple, β flap pale green, β connector vermillion, β protrusion yellow, β fork loop blue, β gate loop orange, βln5 red. β′ subunit elements in Panel C; β′ dock purple, β′ lid yellow, β′ zipper orange and clamp helix (CH) orange, β′ rudder pale green, β′ clamp blue, β′ shelf and jaw dark blue, bridge helix (BH) pink, trigger loop (TL) dark green. Up, and downstream sides of RNAP are indicated for reference.

Transcription initiation involves the RNAP holoenzyme, promoter DNA, and transcription factors (TFs). The primary housekeeping σ transcription factor in *Bacillus subtilis*, σ^A^, contains sub-domains σ_1.1_, σ_2_, σ_3.1_, σ_3.2_, and σ_4_. The holoenzyme structure (core + σ^A^) shown in Fig. 1A is based on the complex with multidrug resistance regulator BmrR (9) and shows an open complex (DNA strands separated at the −10 region, red oval, and template strand inserted in the active site within the primary channel). In this structure much of the lineage-specific βln5 insertion (see below) is missing, but the flexible β-flap tip, important in binding essential transcription factor NusA (15), is visible and binds to the σ_4_ domain that also binds the −35 promoter sequence (red circle, Fig. 1A). In combination with the transcription elongation complex (EC) (10), a near complete structure of the *B. subtilis* enzyme can be modelled. The location of the primary (DNA binding) and secondary (NTP entry) channels are marked as dashed circles in the EC and shown in Fig. 2B. It should be noted that the α-C terminal domain that interacts with both transcription factors and DNA sequences was absent in all RNAP_BS_ structures reported to date, due to the highly flexible sequence connecting the N- and C-terminal domains, and lack of DNA/transcription factor for the α-C terminal domain to bind to. The structure of this important DNA and protein interaction domain has been solved in complex with the transcription factor Spx (16) enabling inclusion in models based on transcription initiation or antitermination complex structures from other organisms (17,18) making it possible to further augment the structures presented here.

**Figure 2.**
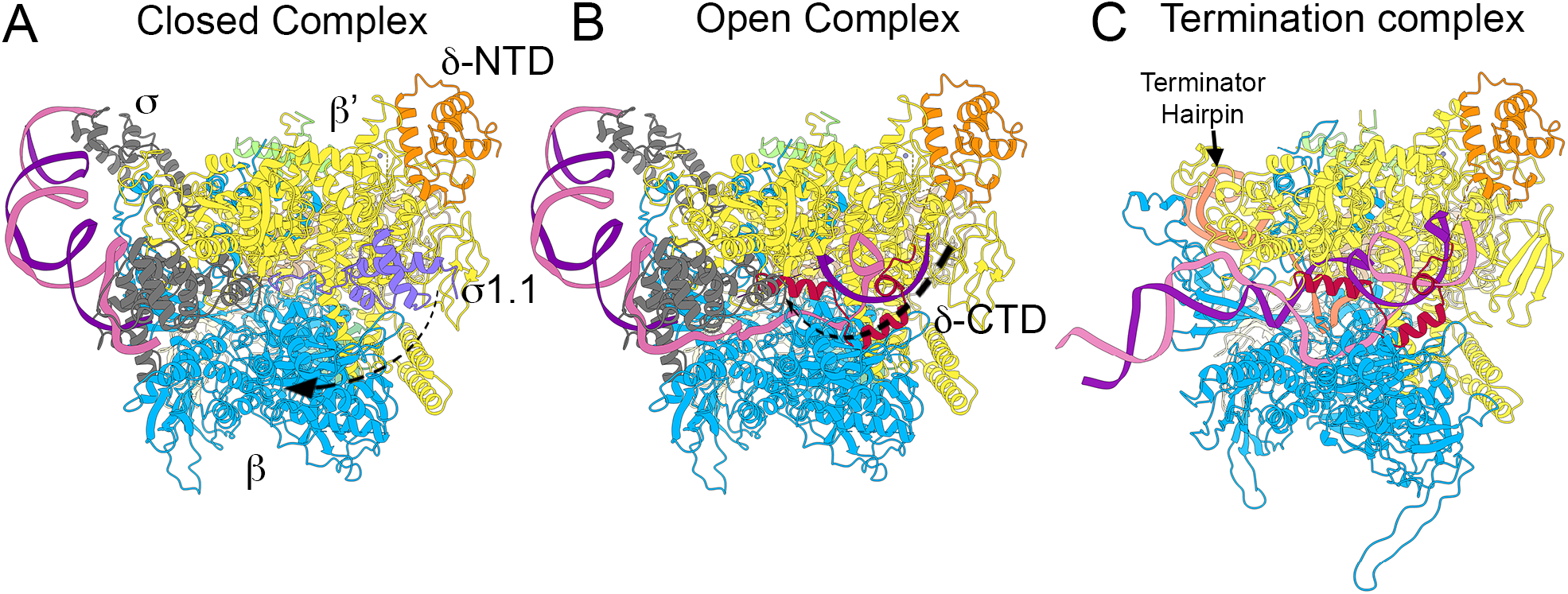
A model for δ subunit activity during transcription initiation and termination. In all panels RNAP subunits are coloured as in Figure 1 with the addition of σ region 1.1 (σ1.1) shown in lavender, the N-terminal domain of δ (δ-NTD) in orange, and the C-terminal domain (δ-CTD) in red. Panel A shows a closed transcription initiation complex with the δ-NTD bound around the β′ shelf. The CTD is not shown (see text for details). σ1.1 is shown in primary channel with the arrow indicating the dissociation of σ1.1 from this site during the transition from a closed to open intiation complex. Panel B shows an open complex in which promoter DNA strands have separated and the template strand has moved into the active site within the primary channel. δ-CTD is shown moving into the primary channel where, due to its polyanionic nature it is able to compete with DNA in the primary channel helping to prevent transcription initiation from cryptic/weak promoters. Panel C shows a model of a transcription termination complex with a RNA hairpin (terminator hairpin).. in the RNA exit channel. The polyanionic δ-CTD is able to disrupt the RNA-DNA hybrid upstream from the active site aiding dissociation of RNA from the complex and RNAP recycling following termination of transcription.

A small, highly conserved, ω subunit binds around the C-terminus of the β′ subunit and is associated with enhancing subunit folding and incorporation of the β′ subunit into the core structure (19,20). The *Bacilli* and *Lactobacilli*, but not *Clostridia*, also possess an additional small subunit named ε, previously annotated as a second ω subunit (dark green, Fig. 1A and B), who’s function has proven to be enigmatic (21–23). Determination of the X-ray crystal structure of ε revealed it showed remarkable similarity to the *E. coli* phage T7 gp2 that binds the β′ jaw region of RNAP (labelled in Fig. 1D) where it inhibits transcription initiation by the host cell RNAP (21,24). Based on this similarity, it was hypothesised that ε could be involved in phage protection through binding to the β′ jaw of RNAP_BS_ (21,23). Determination of the structure of *B. subtilis* holoenzyme and a transcription recycling complex comprising core RNAP with the ATP-dependent remodelling factor HelD revealed ε bound in a pocket formed mainly by the β′ and α_1_ subunits behind the secondary channel on the downstream side of the enzyme (Fig. 1B; (9–11). The location of ε overlaps that of the βln10 and −12 inserts of *Thermus thermophilus* RNAP as well as a region of Archaeal/Eukaryotic Pol II Rpo3/RPB3 associated with enzyme stability, suggesting a similar role for ε in organisms, such as *B. subtilis*, that are meso-thermophilic and capable of vigorous growth up to ~53 °C (10).

Major functional motifs in the β and β′ subunits common to other bacterial RNAPs that are important in DNA strand separation and rewinding (β fork loop, β′-rudder, -lid, -zipper) DNA clamping (β-protrusion, -gate loop, β′ clamp helices, N-terminal β′ clamp), DNA binding cleft flexibility (β switch 3) *etc* are all highly conserved in the *B. subtilis* enzyme and are labelled in Fig. 1C and D (10,11). However, given the differences in aspects of transcription such as open complex stability (above) between *B. subtilis* and *E. coli*, sequence differences and/or insertions (*e.g*. the β′ S13 insertion into the trigger-loop in *E. coli*), these regions remain important areas of focus in structure-function studies in the *Firmicutes*.

Across the eubacteria the β and β′ subunits contain lineage-specific inserts that are largely of unknown function (25,26). Despite the role of most of these lineage-specific inserts being unknown, the βi9 insert in the β subunit of *E. coli* RNAP has recently been putatively implicated in coupling transcription and translation under certain conditions (27). Elucidation of the structure of RNAP_BS_ revealed the structure of the only major lineage-specific insert in *Firmicutes* RNAPs (labelled βln5 in Fig. 1C) (10). Reconciliation of data from previous studies indicates that the βln5 insert is involved in binding to the C-terminal tudor domain of helicase/translocase PcrA (28,29) that is known to interact strongly with RNAP (28,30–32). The βln5 insert is located within the major lobe of the β subunit which is one of the least highly conserved regions in bacterial RNAPs and is the site of many other lineage-specific inserts (25,26) raising the possibility this part of RNAP may be important in providing a platform for lineage-specific transcription factor interaction modules. Overall, due to the lack of lineage-specific inserts (excepting βln5), RNAP from *B. subtilis* and other low G+C Gram positive bacteria represent the smallest multi-subunit RNAPs: *B. subtilis* α_2_ββ′ωε, 352.32 kDa; *Mycobacterium tuberculosis* α_2_ββ′ω, 363.19 kDa; *cf. E. coli* α_2_ββ′ω, 389.05 kDa.

## The δ subunit

Many *Firmicutes* also contain a small δ subunit that is tightly associated with and present at approximately equimolar concentrations with respect to RNAP (30,33). δ is a bipartite protein of 173 amino acids with a globular N-terminal domain and unstructured highly acidic C-terminal domain of approximately equal sizes (34,35). It has been implicated in multiple regulatory roles associated with transcription initiation, inhibition of non-specific transcription, transcription complex recycling and transcription termination (23,36–41). The binding site of δ on RNAP has been the subject of considerable speculation. Independent studies placed it adjacent to the RNA exit channel (42) or close to/inside the DNA binding cleft (43). Determination of the structure of a *B. subtilis* RNAP-HelD-δ recycling complex by (11) showed δ binds on the β′ subunit close to the DNA binding cleft, consistent with findings of *in vivo* cross-linking mass spectrometry studies (43), reconciling the observed biochemical effects of δ on transcription with the structure of a δ-containing transcription complex. In subcellular localisation studies of δ and RNAP in dual fluorescent protein labelled cells, δ perfectly colocalised with RNAP and was present at similar levels suggesting it is associated with RNAP throughout all stages (initiation, elongation, termination) of the transcription cycle (33). Assuming the N-terminal domain of δ remains bound around the β′ jaw/N-terminal β′ clamp region and the acidic unstructured C-terminal domain is both mobile and flexible, we may propose a mechanism for modulation of transcription (Fig. 2). A closed initiation complex, based on the open complex structure of (9) in which the N-terminal σ_1.1_ domain (44) that competes with nucleic acid binding in the primary channel can be modelled *in situ* based on equivalent structures from *E. coli* (24) is shown in Fig. 2A. As the transcription initiation complex transitions from a closed to open complex, the σ_1.1_ domain swings out of the primary channel as unwound DNA enters, placing the single stranded template strand with the transcription start site nucleotide located in position for base pairing with the initiating nucleotide triphosphate (9,45).

Upon σ_1.1_ dissociation during formation of the open complex, the δ C-terminal domain would be able to access DNA within the primary channel (Fig. 2B). The highly negative charge of the δ C-terminal domain would encourage dissociation of weakly bound DNA (poor/non-specific promoter sequences bound by σ-factors) (23,36). The cryo EM structure of the *E. coli* RNAP paused EC (PDB ID 6FLP) was used as a template to model termination hairpin RNA in the RNAP_BS_ EC. The model suggests that the highly flexible C-terminal domain of δ is able to interact with both DNA and the RNA transcript in the active site (Fig. 2B and C), consistent with the observation that δ is much more efficient at displacing RNA from an EC than it is DNA (37). In pause/termination complexes the negatively charged δ C-terminal domain would aid the dissociation of RNA facilitating transcription termination and transcription complex recycling (Fig. 2C) (11,36,41).

## Comparison of RNAP_BS_ with RNAP from other Firmicutes

*B. subtilis* itself is a biologically and industrially significant organism, being important in soil health and promotion of plant growth, protection from plant pathogens, the industrial production of enzymes (*e.g*. proteases/amylases), supplements (*e.g*. nicotinic acid), and antibiotics (*e.g*. bacitracin), as a probiotic, as a foodstuff (*e.g*. in natto), and in the study of regulation of gene expression, especially during cellular differentiation in sporulation (2). As the most studied member of the low G+C Gram-positive *Firmicutes* it is also an important model, and closely related to major pathogens including *B. cereus, B. anthracis, Staphylococcus sp., Streptococcus sp., Enterococcus sp*. and *Clostridium sp*. Organisms such as the *Enterococci* are associated with dissemination of antibiotic resistance determinants, and many clinical isolates of *Staphylococcus* now carry resistance to one or more antibiotics. *S. aureus* is commensal in about 30% of the population and an opportunistic pathogen that remains a major burden on health systems through nosocomial infections, which has been exacerbated in recent years by the rise of community acquired infections (especially methicillin-resistant; MRSA infections) (46). Organisms such as *C. difficile* are associated with diseases difficult to treat successfully with many antibiotics (*e.g. C. difficile* associated diarrhoea (CDAD) relapse is common following vancomycin treatment), and represent a significant burden in terms of both morbidity and mortality to health systems (47). While fidaxomicin was approved by the FDA for treatment of CDAD in 2011, resistance to this drug (lipiarmycin) was first reported in 1977 (48,49), and it is clear new derivatives are needed, as well as a new arsenal of novel compounds to slow the rise of antibiotic resistant infections.

Sequence alignment of RNAP subunits from representatives of these organisms was performed using *B. subtilis* 168, *S. aureus* USA300, *E. faecalis* V583, *S. pyogenes* M1 GAS, *C. difficile* 630 and *C. perfringens* 13 sequences and the resulting CLUSTAL alignment outputs mapped onto the *B. subtilis* elongation complex (PDB ID 6WVJ) in ChimeraX (50,51) with the nucleic acids removed for clarity. The resulting homology sequence maps are shown in Fig. 3 along with a phylogenetic tree produced in MrBayes (52,53) for the *rpoC* (β′ subunit) using the *E. coli rpoC* sequence as an outlier to root the tree. The bootstrap probability values of 1 indicate absolute confidence in the branch divisions and lengths, and agree perfectly, as expected, with the segregation of *B. subtilis* and *S. aureus* to the *Bacilli*, *S. pyogenes* and *E. faecalis* to the *Lactobacilli*, and *C. difficile* and *C. perfringens* to the *Clostridia*.

**Figure 3.**
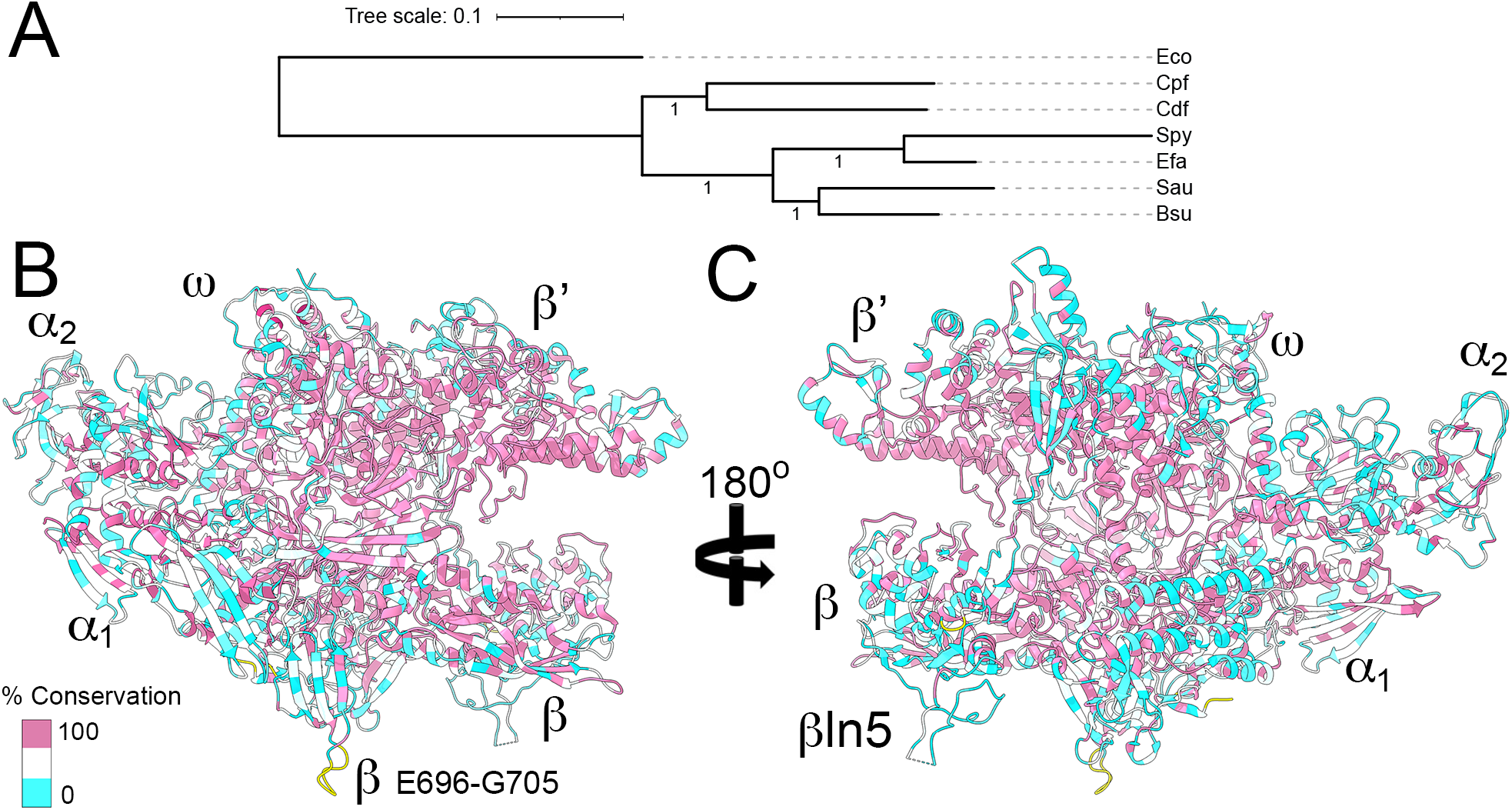
Mapping sequence conservation of pathogenic *Firmicutes* to *B. subtilis* RNAP. Panel A shows a Bayesian tree of sequence alignments of the β subunit of *B. subtilis* (Bsu), *S. aureus* (Sau), *E. faecalis* (Efa), *S. pyogenes* (Spy), *C. difficile* (Cdf), and *C. perfringens* (Cpf). The sequence of *E. coli* β subunit (Eco) was used to root the tree. Tree scale represents amino acid substitutions per site. Bootstrap values are shown on the branches. Panels B and C show up- and downstream views of RNAP, respectively. Subunits are labelled as well as the common to *Firmicutes* βln5 insert, and *B. subtilis*-specific β E696-G705 insert (yellow). A colour scale for sequence conservation shown on the structures is shown on the bottom left of Panel B with sequences 100% conserved pink, 0% conserved cyan and >0, <100 in white.

The level of sequence conservation is high, especially in the β and β′ clamps, active site and β flap where the majority of the functional motifs (see Fig. 1C and D) are found. Although the βln5 is present in all of the organism from which sequences were selected, the level of sequence conservation is relatively low in the major β lobe and βln5 region (Fig. 3C), consistent with the hypothesis (above) that this region maybe important for providing class/order/species-specific binding sites for transcription factors. *B. subtilis*-specific sequences that are absent in the other organisms, such as the 10 amino acid insert at β E696-G705 that protrudes from the bottom of the structure are shown in yellow (Fig. 3B and C). Given the industrial and medical importance of this group of bacteria, regions of identity/difference can be targeted in functional studies or exploited in the rational design of inhibitor compounds as new antimicrobial leads.

## Antimicrobial development options

Transcription is an underutilised target for new antibiotic development, although significant efforts are currently underway to identify promising new leads and to improve the properties of existing clinical compounds (48,54). Many promising compounds fail to make it to market as broad spectrum antibiotics due to the problems of identifying hits that are able to cross the outer membrane of Gram negative bacteria, despite showing excellent activity against Gram positives. Infections due to the *Firmicutes*, including *S. aureus*, *C. difficile*, vancomycin-resistant *Enterococcus*, and drug resistant *Streptococcus* (Group A and B), have been identified by the Centre for Disease Control (CDC) as organisms of major clinical concern for which new approaches/treatments for infection are required (47), and there is a case for developing more narrow spectrum drugs that target this group. Nevertheless, significant hurdles still remain that must be dealt with (55).

Fidaxomicin is a semi-synthetic macrolide that inhibits transcription initiation and was approved for use in CDAD in 2012. *C. difficile* (and other *Clostridia*) are exquisitely sensitive to fidaxomicin, but this is not a property shared by most other *Firmicutes* with some *Streptococci* being > 500 × more resistant than *C. difficile* (56). Structures of fidaxomicin in complex with RNAP from *Mycobacterium tuberculosis* have been solved (57,58) enabling modelling of the drug bound to the *B. subtilis* holoenzyme (PDB ID 7CKQ) with sequence alignments to pathogenic *Firmicutes* homology mapped as in the previous section (Fig. 4A). Other than the homodichloro-orsellenyl moiety that is adjacent to the β-flap tip and −35 promoter sequence binding σ region 4, the bulk of fidaxomicin is buried within the enzyme at the base of the RNAP clamp. All of the RNAP and σ sequences the remaining bulk of fidaxomicin interacts with (switch regions Sw2, Sw3, Sw4 and the σ finger; (57,58)) are highly conserved (pink, Fig. 4A, right side box) consistent with the broad spectrum activity of this compound in *in vitro* transcription assays (although *E. coli* holoenzyme is quite resistant to fidaxomicin; (57)). Increasing the spectrum of activity of fidaxomicin may depend on improving cell permeability properties, especially for organisms such a *S. pneumoniae*, where production of a capsule layer may inhibit efficient cell penetration.

**Figure 4.**
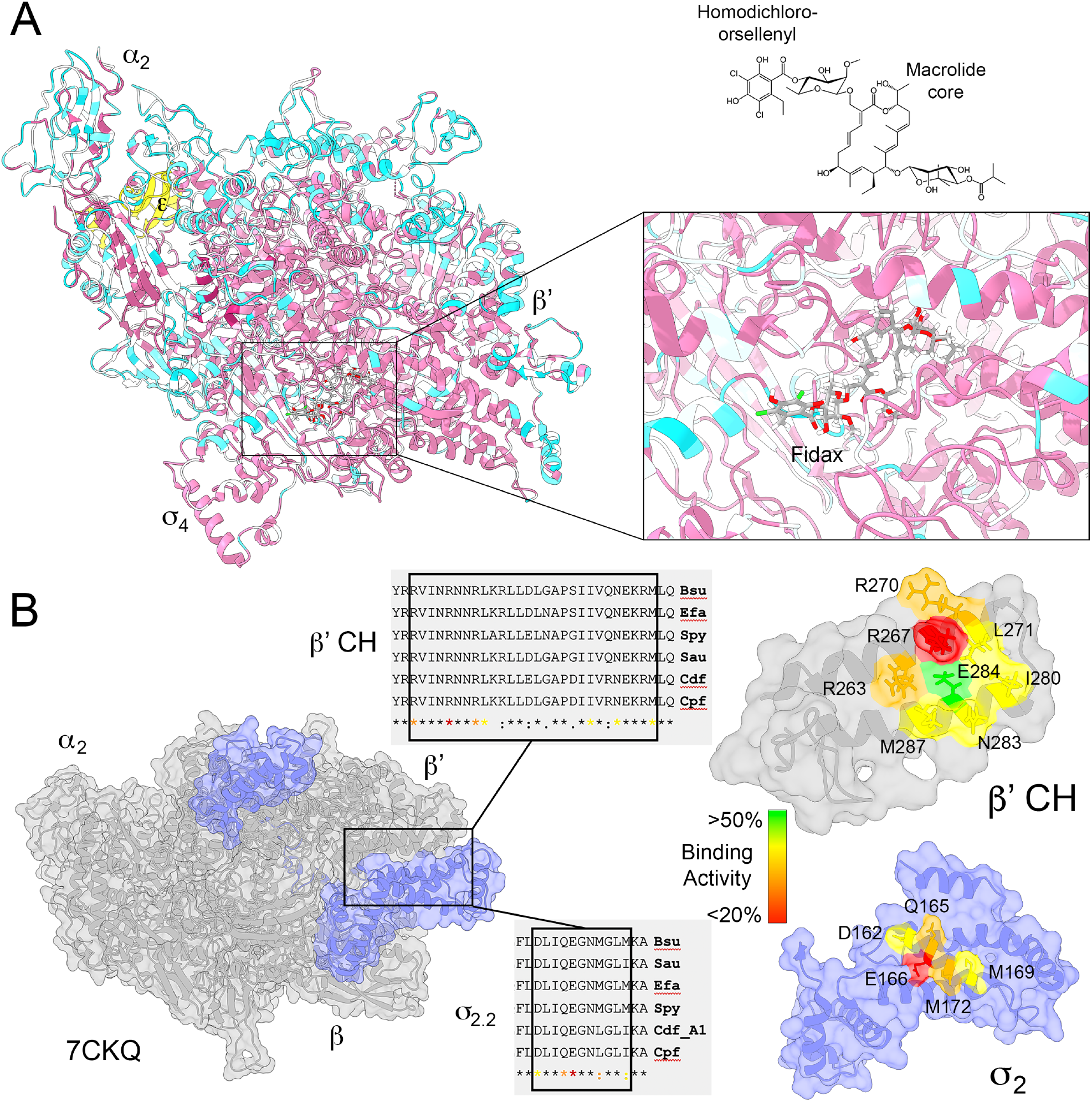
Transcription inhibition drug targets in *Firmicutes* RNAP. Panel A shows sequence conservation mapping of pathogenic *Firmicutes* with colouring as in Figure 3 mapped onto the *B. subtilis* RNAP holoenzyme. The α_2_, and β′ subunits and σ_4_ domain are labelled for reference. The ε subunit which is not conserved in *Clostridia* is shown in yellow. Fidaxomicin (Fidax) is shown docked in the holoenzyme structure (box), which is shown in an enlarged box on the right (see text for details). The structure of fidaxomicin is shown above the right hand box approximately aligned for reference with the homodichloro-orsellenyl and macrolide core labelled for reference. Panel B, left, shows *B. subtilis* holoenzyme with core subunits coloured grey and σ^A^ in pale blue. The box indicates the interaction site between the β′ CH and σ_2.2_ regions essential for formation of holoenzyme. Sequence alignments of the relevant regions are shown adjacent to the holoenzyme. Note, *C. difficile* encodes two σ^A^ subunits, but only the alignment for *sigA1* is shown as *sigA2* is not expressed to a significant level during vegetative growth. Strain labelling is the same as in Figure 3. The right hand side shows enlarged regions of the β′ CH and σ_2.2_ regions with amino acids involved in formation of the holoenzyme colour coded according to their importance as determined from mutagenesis studies (59). The colour ramp indicates the relative binding activity mutation causes to holoenzyme formation.

An area of antimicrobial research that is showing promise for compounds highly active against *Firmicutes* is the development of compounds that inhibit the essential interaction between RNAP and σ^A^. Establishment of a functional holoenzyme complex is dependent on the interaction between a small region of the σ_2.2_ region with the β′ clamp helix (CH) (59,60). These sequences are highly conserved in the *Firmicutes* (Fig. 4B), and across the eubacteria, making this an excellent target for the development of molecules that inhibit this essential protein-protein interaction (PPI) (48,61). Multiple research programs involving high-throughput small molecule screens, structure-based drug design, and small peptide antagonists have yielded promising results (62–71). PPIs are an attractive target for drug development as simultaneous complementary mutations are required at two unlinked genetic loci to confer resistance to a compound whilst retaining the interaction, which has the potential to substantially reduce the rate at which resistance develops (72).

Mutagenesis studies to quantify the importance of specific amino acid residues in formation of holoenzyme have been determined (Fig. 4B right, (59)) that have enabled the construction of pharmacophores for screening compound libraries for potential inhibitor molecules. A major issue with such a target is that the interaction site between σ_2.2_ and the β′ clamp helix is relatively flat making binding specificity and avidity potentially problematic. Nevertheless, compounds have been developed that are highly specific for bacterial initiation complexes, showing no binding activity against human RNAP, and that target initiation complex formation in live bacterial cells as determined by cytological assay (65). Whilst many of these compounds show limited or no activity against Gram negative bacteria, excellent results have been obtained against Gram positive bacteria, including those carrying resistance to multiple antibiotics providing an avenue for development of new drugs effective against Gram positive pathogens (65,66,69,70).

## Concluding statement

Determination of the structure of RNAP from *B. subtilis* now opens the way to undertake detailed structure-function studies on the mechanism of transcription in this Gram-positive model, particularly with respect to mechanistic aspects that are different to *E. coli* helping provide a more holistic understanding of the mechanism of microbial transcription. In addition, this structural information will be important in providing a platform for the rational design and subsequent development of new lead antibiotics to combat infections caused by the *Firmicutes*.

## Acknowledgments

This work was supported by funding from the Australian Research Council (DP210100365) to PJL and AJO. MM was supported by a PhD scholarship from the University of Newcastle Priority Research Centre for Drug Discovery.

## References

1. Browning, D.F. and Busby, S.J. (2016) Local and global regulation of transcription initiation in bacteria. Nat Rev Microbiol, 14, 638–650.

2. Errington, J. and Aart, L.T.V. (2020) Microbe Profile: Bacillus subtilis: model organism for cellular development, and industrial workhorse. Microbiology (Reading), 166, 425–427.

3. Benjin, X. and Ling, L. (2020) Developments, applications, and prospects of cryo-electron microscopy. Protein Sci, 29, 872–882.

4. Chen, J., Wassarman, K.M., Feng, S., Leon, K., Feklistov, A., Winkelman, J.T., Li, Z., Walz, T., Campbell, E.A. and Darst, S.A. (2017) 6S RNA Mimics B-Form DNA to Regulate Escherichia coli RNA Polymerase. Mol Cell, 68, 388–397 e386.

5. Guo, X., Myasnikov, A.G., Chen, J., Crucifix, C., Papai, G., Takacs, M., Schultz, P. and Weixlbaumer, A. (2018) Structural Basis for NusA Stabilized Transcriptional Pausing. Mol Cell, 69, 816–827 e814.

6. Abdelkareem, M., Saint-Andre, C., Takacs, M., Papai, G., Crucifix, C., Guo, X., Ortiz, J. and Weixlbaumer, A. (2019) Structural Basis of Transcription: RNA Polymerase Backtracking and Its Reactivation. Mol Cell, 75, 298–309 e294.

7. Kang, J.Y., Llewellyn, E., Chen, J., Olinares, P.D.B., Brewer, J., Chait, B.T., Campbell, E.A. and Darst, S.A. (2021) Structural basis for transcription complex disruption by the Mfd translocase. Elife, 10.

8. Said, N., Hilal, T., Sunday, N.D., Khatri, A., Burger, J., Mielke, T., Belogurov, G.A., Loll, B., Sen, R., Artsimovitch, I. et al. (2021) Steps toward translocation-independent RNA polymerase inactivation by terminator ATPase rho. Science, 371.

9. Fang, C., Li, L., Zhao, Y., Wu, X., Philips, S.J., You, L., Zhong, M., Shi, X., O’Halloran, T.V., Li, Q. et al. (2020) The bacterial multidrug resistance regulator BmrR distorts promoter DNA to activate transcription. Nat Commun, 11, 6284.

10. Newing, T.P., Oakley, A.J., Miller, M., Dawson, C.J., Brown, S.H.J., Bouwer, J.C., Tolun, G. and Lewis, P.J. (2020) Molecular basis for RNA polymerase-dependent transcription complex recycling by the helicase-like motor protein HelD. Nat Commun, 11, 6420.

11. Pei, H.H., Hilal, T., Chen, Z.A., Huang, Y.H., Gao, Y., Said, N., Loll, B., Rappsilber, J., Belogurov, G.A., Artsimovitch, I. et al. (2020) The delta subunit and NTPase HelD institute a two-pronged mechanism for RNA polymerase recycling. Nat Commun, 11, 6418.

12. Ishikawa, S., Oshima, T., Kurokawa, K., Kusuya, Y. and Ogasawara, N. (2010) RNA polymerase trafficking in Bacillus subtilis cells. J Bacteriol, 192, 5778–5787.

13. Mooney, R.A., Davis, S.E., Peters, J.M., Rowland, J.L., Ansari, A.Z. and Landick, R. (2009) Regulator trafficking on bacterial transcription units in vivo. Mol Cell, 33, 97–108.

14. Krasny, L. and Gourse, R.L. (2004) An alternative strategy for bacterial ribosome synthesis: Bacillus subtilis rRNA transcription regulation. EMBO J, 23, 4473–4483.

15. Ma, C., Mobli, M., Yang, X., Keller, A.N., King, G.F. and Lewis, P.J. (2015) RNA polymerase-induced remodelling of NusA produces a pause enhancement complex. Nucleic Acids Res, 43, 2829–2840.

16. Newberry, K.J., Nakano, S., Zuber, P. and Brennan, R.G. (2005) Crystal structure of the Bacillus subtilis anti-alpha, global transcriptional regulator, Spx, in complex with the alpha C-terminal domain of RNA polymerase. Proc Natl Acad Sci U S A, 102, 15839–15844.

17. Hudson, B.P., Quispe, J., Lara-Gonzalez, S., Kim, Y., Berman, H.M., Arnold, E., Ebright, R.H. and Lawson, C.L. (2009) Three-dimensional EM structure of an intact activator-dependent transcription initiation complex. Proc Natl Acad Sci U S A, 106, 19830–19835.

18. Krupp, F., Said, N., Huang, Y.H., Loll, B., Burger, J., Mielke, T., Spahn, C.M.T. and Wahl, M.C. (2019) Structural Basis for the Action of an All-Purpose Transcription Anti-termination Factor. Mol Cell, 74, 143–157 e145.

19. Ghosh, P., Ishihama, A. and Chatterji, D. (2001) Escherichia coli RNA polymerase subunit omega and its N-terminal domain bind full-length beta’ to facilitate incorporation into the alpha2beta subassembly. Eur J Biochem, 268, 4621–4627.

20. Minakhin, L., Bhagat, S., Brunning, A., Campbell, E.A., Darst, S.A., Ebright, R.H. and Severinov, K. (2001) Bacterial RNA polymerase subunit omega and eukaryotic RNA polymerase subunit RPB6 are sequence, structural, and functional homologs and promote RNA polymerase assembly. Proc Natl Acad Sci U S A, 98, 892–897.

21. Keller, A.N., Yang, X., Wiedermannova, J., Delumeau, O., Krasny, L. and Lewis, P.J. (2014) epsilon, a new subunit of RNA polymerase found in gram-positive bacteria. J Bacteriol, 196, 3622–3632.

22. Spiegelman, G.B., Hiatt, W.R. and Whiteley, H.R. (1978) Role of the 21,000 molecular weight polypeptide of Bacillus subtilis RNA polymerase in RNA synthesis. J Biol Chem, 253, 1756–1765.

23. Weiss, A. and Shaw, L.N. (2015) Small things considered: the small accessory subunits of RNA polymerase in Gram-positive bacteria. FEMS Microbiol Rev, 39, 541–554.

24. Bae, B., Davis, E., Brown, D., Campbell, E.A., Wigneshweraraj, S. and Darst, S.A. (2013) Phage T7 Gp2 inhibition of Escherichia coli RNA polymerase involves misappropriation of sigma70 domain 1.1. Proc Natl Acad Sci U S A, 110, 19772–19777.

25. Lane, W.J. and Darst, S.A. (2010) Molecular evolution of multisubunit RNA polymerases: structural analysis. J Mol Biol, 395, 686–704.

26. Lane, W.J. and Darst, S.A. (2010) Molecular evolution of multisubunit RNA polymerases: sequence analysis. J Mol Biol, 395, 671–685.

27. Wang, C., Molodtsov, V., Firlar, E., Kaelber, J.T., Blaha, G., Su, M. and Ebright, R.H. (2020) Structural basis of transcription-translation coupling. Science, 369, 1359–1365.

28. Harriott, K. (2012), The University of Newcaslte.

29. Sanders, K., Lin, C.L., Smith, A.J., Cronin, N., Fisher, G., Eftychidis, V., McGlynn, P., Savery, N.J., Wigley, D.B. and Dillingham, M.S. (2017) The structure and function of an RNA polymerase interaction domain in the PcrA/UvrD helicase. Nucleic Acids Res, 45, 3875–3887.

30. Delumeau, O., Lecointe, F., Muntel, J., Guillot, A., Guedon, E., Monnet, V., Hecker, M., Becher, D., Polard, P. and Noirot, P. (2011) The dynamic protein partnership of RNA polymerase in Bacillus subtilis. Proteomics, 11, 2992–3001.

31. Gwynn, E.J., Smith, A.J., Guy, C.P., Savery, N.J., McGlynn, P. and Dillingham, M.S. (2013) The conserved C-terminus of the PcrA/UvrD helicase interacts directly with RNA polymerase. PLoS One, 8, e78141.

32. Noirot-Gros, M.F., Dervyn, E., Wu, L.J., Mervelet, P., Errington, J., Ehrlich, S.D. and Noirot, P. (2002) An expanded view of bacterial DNA replication. Proc Natl Acad Sci U S A, 99, 8342–8347.

33. Doherty, G.P., Fogg, M.J., Wilkinson, A.J. and Lewis, P.J. (2010) Small subunits of RNA polymerase: localization, levels and implications for core enzyme composition. Microbiology, 156, 3532–3543.

34. Kuban, V., Srb, P., Stegnerova, H., Padrta, P., Zachrdla, M., Jasenakova, Z., Sanderova, H., Vitovska, D., Krasny, L., Koval, T. et al. (2019) Quantitative Conformational Analysis of Functionally Important Electrostatic Interactions in the Intrinsically Disordered Region of Delta Subunit of Bacterial RNA Polymerase. J Am Chem Soc, 141, 16817–16828.

35. Motackova, V., Sanderova, H., Zidek, L., Novacek, J., Padrta, P., Svenkova, A., Korelusova, J., Jonak, J., Krasny, L. and Sklenar, V. (2010) Solution structure of the N-terminal domain of Bacillus subtilis delta subunit of RNA polymerase and its classification based on structural homologs. Proteins, 78, 1807–1810.

36. Juang, Y.L. and Helmann, J.D. (1994) The delta subunit of Bacillus subtilis RNA polymerase. An allosteric effector of the initiation and core-recycling phases of transcription. J Mol Biol, 239, 1–14.

37. Lopez de Saro, F.J., Woody, A.Y. and Helmann, J.D. (1995) Structural analysis of the Bacillus subtilis delta factor: a protein polyanion which displaces RNA from RNA polymerase. J Mol Biol, 252, 189–202.

38. Lopez de Saro, F.J., Yoshikawa, N. and Helmann, J.D. (1999) Expression, abundance, and RNA polymerase binding properties of the delta factor of Bacillus subtilis. J Biol Chem, 274, 15953–15958.

39. Nicolas, P., Mader, U., Dervyn, E., Rochat, T., Leduc, A., Pigeonneau, N., Bidnenko, E., Marchadier, E., Hoebeke, M., Aymerich, S. et al. (2012) Condition-dependent transcriptome reveals high-level regulatory architecture in Bacillus subtilis. Science, 335, 1103–1106.

40. Rabatinova, A., Sanderova, H., Jirat Matejckova, J., Korelusova, J., Sojka, L., Barvik, I., Papouskova, V., Sklenar, V., Zidek, L. and Krasny, L. (2013) The delta subunit of RNA polymerase is required for rapid changes in gene expression and competitive fitness of the cell. J Bacteriol, 195, 2603–2611.

41. Wiedermannova, J., Sudzinova, P., Koval, T., Rabatinova, A., Sanderova, H., Ramaniuk, O., Rittich, S., Dohnalek, J., Fu, Z., Halada, P. et al. (2014) Characterization of HelD, an interacting partner of RNA polymerase from Bacillus subtilis. Nucleic Acids Res, 42, 5151–5163.

42. Prajapati, R.K., Sengupta, S., Rudra, P. and Mukhopadhyay, J. (2016) Bacillus subtilis delta Factor Functions as a Transcriptional Regulator by Facilitating the Open Complex Formation. J Biol Chem, 291, 1064–1075.

43. de Jong, L., de Koning, E.A., Roseboom, W., Buncherd, H., Wanner, M.J., Dapic, I., Jansen, P.J., van Maarseveen, J.H., Corthals, G.L., Lewis, P.J. et al. (2017) In-Culture Cross-Linking of Bacterial Cells Reveals Large-Scale Dynamic Protein-Protein Interactions at the Peptide Level. J Proteome Res, 16, 2457–2471.

44. Zachrdla, M., Padrta, P., Rabatinova, A., Sanderova, H., Barvik, I., Krasny, L. and Zidek, L. (2017) Solution structure of domain 1.1 of the sigma(A) factor from Bacillus subtilis is preformed for binding to the RNA polymerase core. J Biol Chem, 292, 11610–11617.

45. Murakami, K.S. and Darst, S.A. (2003) Bacterial RNA polymerases: the wholo story. Curr Opin Struct Biol, 13, 31–39.

46. Tong, S.Y., Davis, J.S., Eichenberger, E., Holland, T.L. and Fowler, V.G., Jr. (2015) Staphylococcus aureus infections: epidemiology, pathophysiology, clinical manifestations, and management. Clin Microbiol Rev, 28, 603–661.

47. CDC. (2013), Centre for Disease Control.

48. Ma, C., Yang, X. and Lewis, P.J. (2016) Bacterial Transcription as a Target for Antibacterial Drug Development. Microbiol Mol Biol Rev, 80, 139–160.

49. Sonenshein, A.L., Alexander, H.B., Rothstein, D.M. and Fisher, S.H. (1977) Lipiarmycin-resistant ribonucleic acid polymerase mutants of Bacillus subtilis. J Bacteriol, 132, 73–79.

50. Goddard, T.D., Huang, C.C., Meng, E.C., Pettersen, E.F., Couch, G.S., Morris, J.H. and Ferrin, T.E. (2018) UCSF ChimeraX: Meeting modern challenges in visualization and analysis. Protein Sci, 27, 14–25.

51. Pettersen, E.F., Goddard, T.D., Huang, C.C., Meng, E.C., Couch, G.S., Croll, T.I., Morris, J.H. and Ferrin, T.E. (2020) UCSF ChimeraX: Structure Visualization for Researchers, Educators, and Developers. Protein Sci.

52. Huelsenbeck, J.P. and Ronquist, F. (2001) MRBAYES: Bayesian inference of phylogenetic trees. Bioinformatics, 17, 754–755.

53. Ronquist, F., Teslenko, M., van der Mark, P., Ayres, D.L., Darling, A., Hohna, S., Larget, B., Liu, L., Suchard, M.A. and Huelsenbeck, J.P. (2012) MrBayes 3.2: efficient Bayesian phylogenetic inference and model choice across a large model space. Syst Biol, 61, 539–542.

54. Maffioli, S.I., Zhang, Y., Degen, D., Carzaniga, T., Del Gatto, G., Serina, S., Monciardini, P., Mazzetti, C., Guglierame, P., Candiani, G. et al. (2017) Antibacterial Nucleoside-Analog Inhibitor of Bacterial RNA Polymerase. Cell, 169, 1240–1248 e1223.

55. Alm, R.A. and Lahiri, S.D. (2020) Narrow-Spectrum Antibacterial Agents-Benefits and Challenges. Antibiotics (Basel), 9.

56. Goldstein, E.J., Babakhani, F. and Citron, D.M. (2012) Antimicrobial activities of fidaxomicin. Clin Infect Dis, 55 Suppl 2, S143–148.

57. Boyaci, H., Chen, J., Lilic, M., Palka, M., Mooney, R.A., Landick, R., Darst, S.A. and Campbell, E.A. (2018) Fidaxomicin jams Mycobacterium tuberculosis RNA polymerase motions needed for initiation via RbpA contacts. Elife, 7.

58. Lin, W., Das, K., Degen, D., Mazumder, A., Duchi, D., Wang, D., Ebright, Y.W., Ebright, R.Y., Sineva, E., Gigliotti, M. et al. (2018) Structural Basis of Transcription Inhibition by Fidaxomicin (Lipiarmycin A3). Mol Cell, 70, 60–71 e15.

59. Johnston, E.B., Lewis, P.J. and Griffith, R. (2009) The interaction of Bacillus subtilis sigmaA with RNA polymerase. Protein Sci, 18, 2287–2297.

60. Arthur, T.M., Anthony, L.C. and Burgess, R.R. (2000) Mutational analysis of beta ‘260-309, a sigma 70 binding site located on Escherichia coli core RNA polymerase. J Biol Chem, 275, 23113–23119.

61. Cossar, P.J., Lewis, P.J. and McCluskey, A. (2020) Protein-protein interactions as antibiotic targets: A medicinal chemistry perspective. Med Res Rev, 40, 469–494.

62. Andre, E., Bastide, L., Michaux-Charachon, S., Gouby, A., Villain-Guillot, P., Latouche, J., Bouchet, A., Gualtieri, M. and Leonetti, J.P. (2006) Novel synthetic molecules targeting the bacterial RNA polymerase assembly. J Antimicrob Chemother, 57, 245–251.

63. Glaser, B.T., Bergendahl, V., Thompson, N.E., Olson, B. and Burgess, R.R. (2007) LRET-based HTS of a small-compound library for inhibitors of bacterial RNA polymerase. Assay Drug Dev Technol, 5, 759–768.

64. Husecken, K., Negri, M., Fruth, M., Boettcher, S., Hartmann, R.W. and Haupenthal, J. (2013) Peptide-Based Investigation of the Escherichia coli RNA Polymerase sigma(70):Core Interface As Target Site. ACS Chem Biol.

65. Ma, C., Yang, X. and Lewis, P.J. (2016) Bacterial Transcription Inhibitor of RNA Polymerase Holoenzyme Formation by Structure-Based Drug Design: From in Silico Screening to Validation. ACS Infect Dis, 2, 39–46.

66. Ma, C., Yang, X., Kandemir, H., Mielczarek, M., Johnston, E.B., Griffith, R., Kumar, N. and Lewis, P.J. (2013) Inhibitors of bacterial transcription initiation complex formation. ACS Chem Biol, 8, 1972–1980.

67. Mielczarek, M., Thomas, R.V., Ma, C., Kandemir, H., Yang, X., Bhadbhade, M., Black, D.S., Griffith, R., Lewis, P.J. and Kumar, N. (2015) Synthesis and biological activity of novel mono-indole and mono-benzofuran inhibitors of bacterial transcription initiation complex formation. Bioorg Med Chem, 23, 1763–1775.

68. Wenholz, D.S., Zeng, M., Ma, C., Mielczarek, M., Yang, X., Bhadbhade, M., Black, D.S.C., Lewis, P.J., Griffith, R. and Kumar, N. (2017) Small molecule inhibitors of bacterial transcription complex formation. Bioorg Med Chem Lett, 27, 4302–4308.

69. Ye, J., Chu, A.J., Harper, R., Chan, S.T., Shek, T.L., Zhang, Y., Ip, M., Sambir, M., Artsimovitch, I., Zuo, Z. et al. (2020) Discovery of Antibacterials That Inhibit Bacterial RNA Polymerase Interactions with Sigma Factors. J Med Chem, 63, 7695–7720.

70. Ye, J., Chu, A.J., Lin, L., Chan, S.T., Harper, R., Xiao, M., Artsimovitch, I., Zuo, Z., Ma, C. and Yang, X. (2020) Benzyl and benzoyl benzoic acid inhibitors of bacterial RNA polymerase-sigma factor interaction. Eur J Med Chem, 208, 112671.

71. Sartini, S., Levati, E., Maccesi, M., Guerra, M., Spadoni, G., Bach, S., Benincasa, M., Scocchi, M., Ottonello, S., Rivara, S. et al. (2019) New Antimicrobials Targeting Bacterial RNA Polymerase Holoenzyme Assembly Identified with an in Vivo BRET-Based Discovery Platform. ACS Chem Biol, 14, 1727–1736.

72. Igler, C., Rolff, J. and Regoes, R. (2021) Multi-step vs. single-step resistance evolution under different drugs, pharmacokinetics and treatment regimens. Elife, 10.

